# Behavioral Opportunism and Altered Dopamine Dynamics in Mice Exposed to Early Life Adversity

**DOI:** 10.64898/2025.12.03.692237

**Authors:** Meghan E. Gallo, Arif A. Hamid, Alana Jaskir, Tracy Pan, Dayshalis Ofray, Camila DeMaestri, Jocelyn Breton, A. Ian More, Michael J. Frank, Christopher I. Moore, Kevin G. Bath

## Abstract

Early life adversity (ELA) confers risk for reward-related psychopathologies. These risks may stem from adaptations optimizing reward pursuit in anticipation of unreliable, resource poor environments. One rational adaptation to poor, unreliable environments is Behavioral Opportunism: updating expectations more slowly and acting vigorously only when reward is immediately available. To systematically test the impact of ELA on behavioral strategies and underlying reward processing mechanisms, we exposed mice to resource restriction (limited bedding and nesting materials for 7 days) to manipulate the reliability and quality of early life care. Subsequently, we tested adults’ reward learning and decision making in a two-arm bandit task and recorded dopamine signaling using dLight1.2 fiber photometry in the nucleus accumbens core. Exposure to ELA led to poorer choice discrimination, impaired learning, and decreased adaptation to changes in reward availability. Furthermore, ELA mice were slower to choose between levers but were faster to retrieve immediately available rewards when delivered, consistent with a strategy of behavioral opportunism. Dopamine signaling predicted behavior in both rearing conditions, and its fluctuations were strongly predictive of faster retrieval in ELA mice and an increased likelihood of choice repetition, implying that aberrant dopamine signals underlie slowed learning and vigorous action for immediately available rewards. To understand key features of maternal interactions driving these effects, we used home cage video monitoring to quantify maternal behaviors, continuously, across early life. We found that specific experiential outcomes, such as maternal kicking, intensified behavioral opportunism in adults, predicting poorer bandit task performance beyond the group effect of ELA. Behavioral opportunism provides an explanatory framework for interpreting altered reward processing and reward pursuit in adulthood for individuals exposed to ELA.

## Introduction

Early life adversity (ELA) is associated with increased risk for reward-related psychopathologies, including substance use disorder, depression, schizophrenia, and bipolar disorder^1–6^. Increased psychopathology risk may reflect neural and behavioral adaptations optimizing reward prediction and pursuit in adverse early environments, including resource scarcity, neglect, or abuse.

For example, a behavioral adaptation to resource-scarce environments entails updating expectations more slowly, to marginalize over volatile outcomes, discounting distant rewards to favor proximal options, and acting opportunistically when rewards are immediately available. These differences, which we refer to as *Behavioral Opportunism*, may be adaptive in response to specific forms of adverse early environments, such as resource scarcity and unreliability, but then persist beyond them. Indeed, strategies learned early in development may impact lifelong behavior. Humans exposed to ELA exhibit these tendencies later in life: reacting more strongly to salient and immediately available rewards^7–9^, discounting delayed rewards more steeply^10–14^, and learning more slowly^15,16^. These behavioral patterns imply risk for reward-related psychopathologies^17^. It is unclear, however, how ELA alters the mechanisms driving behavioral opportunism.

Striatal dopamine likely contributes to these ELA-related behavioral changes. Dopamine signaling plays well-established roles in promoting plasticity at cortico-striatal synapses, thereby driving reward learning^18,19^ . A rich literature also links dopamine to reward sensitivity^20–23^, reward discounting^24,25^, and willingness to expend effort for reward^26–31^. We thus hypothesize that ELA may alter the dopamine system driving both slowed learning and motivation for immediately available rewards.

Striatal dopamine signaling has been implicated in diverse temporal dynamics with dissociable consequences for learning, and motivation. Phasic and tonic activity of midbrain dopamine cells^32,33^ have been linked, respectively, with learning and performance. This suggests dual behavioral roles of midbrain dopamine neural activity, yet more recent data paint a richer picture of the role of dopamine dynamics^27,34,35^. Key work using cyclic voltammetry during a two-arm probabilistic reward task revealed that striatal dopamine tone encodes both the temporally-discounted value of future rewards, and reward history – increasing as animals get closer to rewards and when the recent environment delivered rewards more frequently^27^. In sum, findings indicate that striatal dopamine tone encodes the expected value of the environment (‘state value’ in reinforcement learning terms) and regulates willingness to work, dynamically^27,34,35^. These observations suggest that dopamine dynamics may drive dynamic changes in vigor, as a function of reward proximity, in behavioral opportunism.

Human studies reveal altered reward processing among adults who have experienced ELA, consistent with patterns of behavioral opportunism. In line with the prediction that adults exposed to ELA are more responsive to immediately available rewards and less responsive to distant rewards, Boecker et al. demonstrated increased activation of reward-processing brain regions including the insula, right palladium and bilateral putamen, in response to cues predicting a certain reward and decreased activation of ventral striatum, putamen and thalamus in anticipation of uncertain rewards^36^. Similarly, experiences of ELA predict blunted subjective responses to anticipatory cues of uncertain rewards as indexed by aberrant left basal ganglia activity^1^. Differences in reward processing may reflect the impact of ELA on dopamine synthesis capacity, which is elevated as measured by 18F-DOPA positron emission tomography^37^. However, beyond showing that DA signaling and reward processing are altered, these studies raise the question: how does ELA alter the dynamic regulation of learning and decision making by DA signaling?

Here, we test the hypothesis that animals exposed to ELA adopt a strategy of behavioral opportunism – adapting more slowly to changes in environmental reward statistics and retrieving immediately available rewards more vigorously – and that these behavioral patterns persist into adulthood. We further hypothesize that aberrant patterns of striatal dopamine signaling mediate these effects of ELA on reward learning and decision-making.

## Results

We used a two-armed probabilistic bandit task to evaluate the effects of early life adversity (ELA; using a limited and bedding manipulation) on adult learning and decision making, while systematically manipulating reward availability/richness (**Figure 1A**). In adulthood, animals reared in control (n = 38) and ELA conditions (n = 39) were presented with two levers, on each trial, with differing reward probabilities. The probabilities of reward payout for each lever varied across four block types: easy discrimination with 10 versus 90% payout (10:90), reward rich blocks with 50 and 90% (50:90), reward poor blocks with 10 and 50% (10:50), and chance-level blocks with 50 and 50% (50:50). Reward probabilities pseudo-randomly alternated across blocks and the order was left-right counterbalanced across sessions. Reward rates switched randomly after 25-35 trials in each block, forcing animals to continuously sample both levers, to mazimize reward rates across blocks.

**Figure 1:**
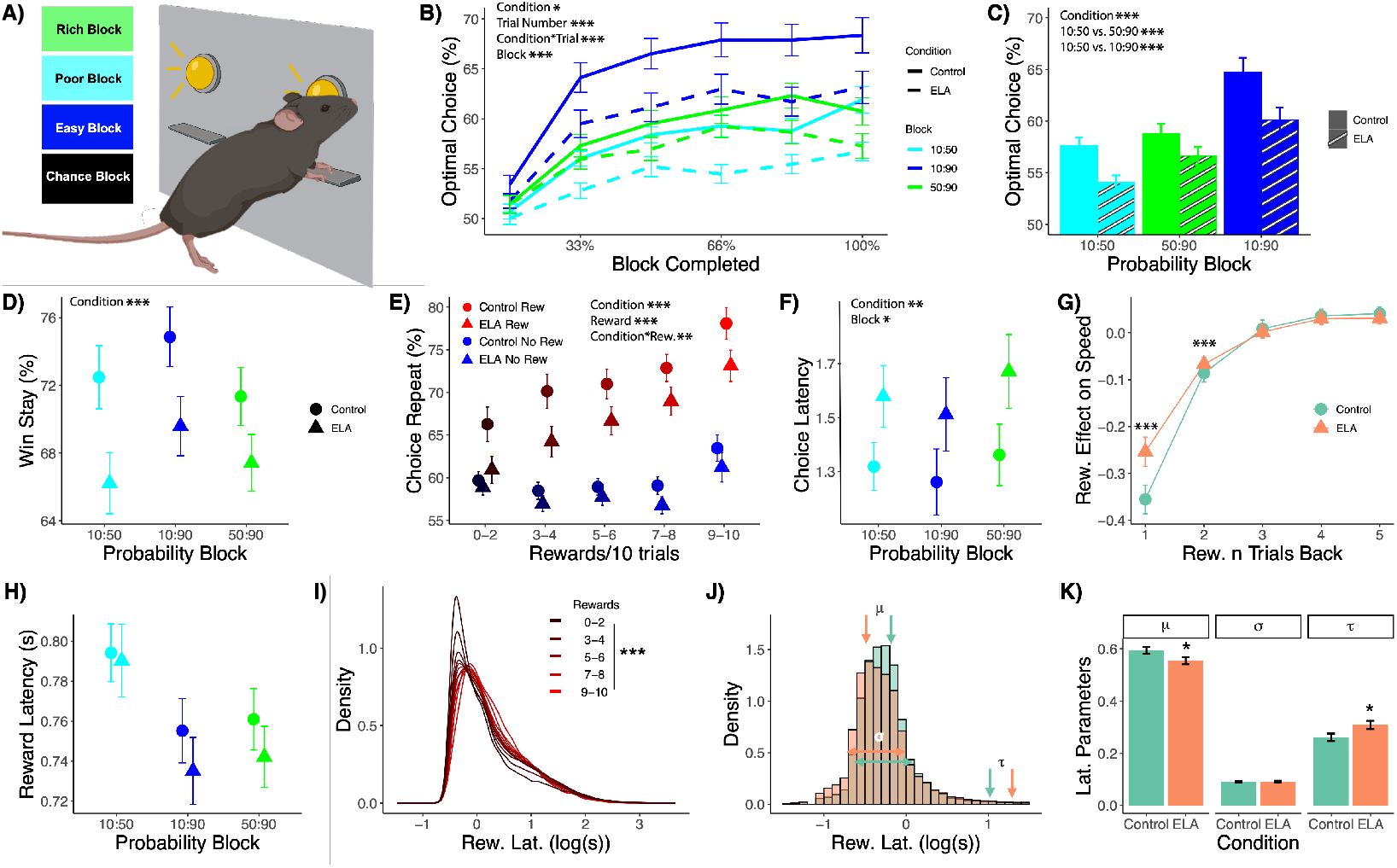
Bandit Task Behavior. **(A)** On each trial, ELA and Control animals chose between two levers that vary in reward probability every 25-35 trials. Rich blocks have a payout rate of 90% for the optimal lever and 50% for the suboptimal, poor-50% and 10%, easy discrimination-90% and 10%, and chance-50% for both levers. **(B)** The rate at which animals chose the optimal lever – defined as the lever with the highest reward probability payout in each block – increases as animals learn reward contingencies through trial and error. Optimal choice rates vary by block and group, with faster learning in easy discrimination blocks versus rich blocks versus poor blocks, and Control animals learning faster than ELA animals. **(C)** Asymptotic optimal choice rates (proportion of optimal choices in the last half of every block) also vary by block and group with best performance in the easy versus the rich and versus the poor blocks and Control versus ELA animals. **(D)** ELA animals win and stay with previous choice less than controls. **(E)** Rates of repeating a choice are higher after a reward versus no reward and vary by reward history. Animals are more likely to repeat rewarded actions if they experienced more rewards on the last 10 trials (regardless of action taken) and unrewarded actions if there were 9 or 10 rewards on the last 10 trials. **(F)** The median time to press a lever after options are presented is slower for ELA versus Control animals, across all blocks. **(G)** Rewards *n* trials back speed choices on the current trial, with the biggest effect for rewards 1 trial back. The speeding effects at 1 and 2 trials back are smaller for ELA versus control animals. **(H)** Median latency to retrieve rewards across blocks. **(I)** Reaction time distributions show a clear, parametric effect of reward history with faster choices when there were more rewards in the last 10 trials. **(J)** Histogram of latency to retrieve rewards reveals a left shifted distribution in ELA animals. **(K)** An ex-Gaussian decomposition of reaction times reveals that ELA animals have a reliably faster expectation for reward retrieval, and also have a bigger tail, indicating a greater proportion of very long retrieval times. *All error bars and bands are +/- SEM*.

### Early life adversity impairs learning and decision making

The rate of optimal (higher probability) lever choices increased across trials in each block, scaled with the block type^27,38^, and differed by condition (**Figure 1B**). As predicted, a mixed-effects regression of trial-wise optimal choice on block type, trial number, condition, and their interaction, while controlling for sex, revealed that animals chose the higher probability lever more often in the easy discrimination block versus the poor block (β = 0.311, *p* < 2.00 ×10^-16^) and in the rich block versus the poor block (β = 0.0578, p = 0.0111). Better performance in the rich versus poor block, despite an equivalent difference in the reward rates, was predicted based on the hypothesized capacity of striatal dopamine signaling to enhance discrimination between good versus better options in rich environments^39^. Compared to controls, ELA blunts performance across all three blocks (β = -0.144, *p* = 3.77 × 10^-4^), while preserving performance effects by block as revealed by a generalized logistic regression. Our regression also revealed a main effect of trial number within a block (β = 0.180, *p* < 2.00×10^-16^), and an interaction of condition and trial number (β = -0.0594, *p* = 0.0166). The significant interaction demonstrates that ELA animals also learn at a slower rate across trials, within a block.

We also evaluated asymptotic performance, as defined by optimal choice in the last half of trials in each block, with a generalized, mixed-effects regression of optimal choice on block type, condition, and their interaction, while controlling for sex. Animals perform better in the easy discrimination block versus the poor block (β = 0.369, *p* < 2.00 ×10^-16^) and in the rich block versus the poor block (β = 0.0667, p = 0.0304). Critically, ELA animals choose the optimal, higher reward probability lever at a lower rate (β = -0.182, *p* = 3.72 ×10^-4^). Sex also influenced performance, with females choosing optimally at a lower rate, asymptotically (β = -0.0926, *p* = 0.0354). There were no interactions between condition and block type (p’s > 0.264), indicating a main effect of exposure to adverse rearing on asymptotic choice only.

We also investigated the immediate effect of reward outcomes on behavior across groups (**Figure 1D**). We fit a generalized, mixed-effects regression of choice repetition following a reward (i.e., “win-stay”) on block type, condition, and their interaction. Animals repeated rewarded choices more often in the easy discrimination block compared to the poor block (β = 0.120, *p* = 6.14× 10^-6^), but less often in the rich block compared to the poor block (β =-0.0783, *p* = 0.00803). Furthermore, ELA animals are less likely to repeat a rewarded choice compared to controls (β = -0.321, *p* = 0.00814). Recent reward history is hypothesized to boost sensitivity to rewards and enhance discrimination^39^. Therefore, we examined the effect of reward history on the probability of repeating the same choice as a function of rewards in the last ten trials (**Figure 1E**). A generalized, mixed-effects logistic regression of choice repetition on trial outcome, condition, reward history, and their interaction indicates that all animals are more likely to repeat a choice immediately following a reward (Intercept: β = 0.318, p < 2.00 × 10^-16^). This effect is compounded if there are more rewards in recent history as indexed by a significant interaction with the linear effect of the number of rewards on the last 10 trials (β = 0.123, p < 2.00 × 10^-16^). Finally, ELA animals are less likely to repeat following a reward (β = - 0.142, p = 0.0366). However, there is no further interaction between condition and reward history (β = 0.00104, p = 0.8827).

While ELA animals are less likely to repeat their choice following a reward, they are also less likely to repeat a choice following a reward omission (β = -0.142, p = 0.0366). However, they are still more likely than chance to repeat choices after reward omissions, reflecting perseverative tendencies (β = 0.683, p = 2 × 10^-16^). Taken together, ELA animals are less likely to repeat a rewarded action and are less likely to select the optimal lever, overall, suggesting a disruption in reinforcement mechanisms.

### ELA animals are slower to choose but faster to seize available rewards

Previous studies in humans demonstrated that ELA is associated with slower reaction times for uncertain rewards^1,36^. Indeed, exposure to ELA slowed choices (**Figure 1F**). A regression of median reaction times by animal on condition and probability block, controlling for sex, revealed that ELA animals are slower across all blocks (β = 0.398, p = 0.00922), and that all animals respond more slowly in the 50:50 (chance) block in contrast with the poor block (β = 0.0551, p = 0.0332). Thus, as expected, all animals are slower to choose when options are harder to discriminate at chance, and ELA animals are overall slower to choose.

We next examined whether animals are faster to choose when they are in a locally rich environment as defined by having received more rewards in recent history. Prior work has shown behavioral invigoration in locally rich environments^27^. We find that reward history impacts choice latency as a function of trial number over the last five trials (**Figure 1G**). Both ELA animals and control animals show the biggest effect of reward one trial back on latency to choose (β = -0.350, p = 3.65 × 10^-14^) and a smaller effect of reward received two trials back (β = -0.109, p < 2.00 × 10^-16^). Importantly, we also observed a condition by reward history interaction: in ELA animals, the effects of reward one trial back (β = 9.65× 10^-2^, p = 0.0495) and two trials back (β = 0.0326, p = 1.61 × 10^-5^) to speed reward retrieval is smaller among ELA animals than the effect in control animals. That is, while recent rewards speed responding on the subsequent trial for both groups, this effect is blunted among ELA animals, again supporting the hypothesis that ELA animals are slower to update their expectations across reward outcomes.

Latency to retrieve reward is also impacted by group, probability block, and reward history. There is an effect of block on reward retrival latency, as animals in both groups are faster to retrieve rewards in the rich (β = -0.350, p = 5.70 × 10^-11^) and easy discrimation (β = -0.489, p = 4.53 × 10^-8^) blocks compared to the poor block (**Figure 1H**). Futhermore, histograms of reaction times reveal a clear parametric effect of reward history on the latency to retrieve, with faster reaction times following more rewards in the last 10 trials (**Figure 1I**; β = -0.0294, p = 1.82×10^-6^).

In contrast to slower choice reaction times, ELA animals retrieve immediately available rewards faster than controls (**Figure 1J-K**). An ex-Gaussian analysis of latency to retrieve rewards was used to dissect reaction time distributions into parameters reflecting the central tendency (*μ*), variance (*σ*), and skew (*τ*). These analyses revealed that ELA animals are faster to retrieve rewards, on average (*µ*, t = -2.01, p = .0498). The analysis also revealed that they are less consistent than controls with a higher proportion of very slow retrieval trials (*τ*, t = 2.14 p = .0365) – potentially reflecting a higher proportion of disengaged trials.

ELA thus alters bandit task performance in three key ways that are consistent with behavioral opportunism: 1) slower updating of reward expectations, 2) slower action when rewards are uncertain, and 3) faster action when rewards are immediately available. As dopamine signaling has been implicated in dissociable effects on both the learning and execution of actions, we next assessed the effects dopamine on task performance and how that relationship is altered by ELA.

### Dopamine release reflects rewards and punishments, and tracks reward history

For a subset of animals in both groups, we next recorded dopamine release in the nucleus accumbens core during the probabilistic bandit task using dLight1.2 fiber photometry (**Figure 2A**). dLight signal was high-pass filtered and z-scored within-session, prior to analyses. As expected, there is a phasic increase in dLight signal when animals receive rewards and a dip when rewards are omitted, consistent with the reward prediction error hypothesis of phasic dopamine release (**Figure 2B-E**). Moreover, there is a clear parametric effect of reward history (defined as the cumulative number of rewards on the past 10 trials) such that phasic dopamine release is greater when there are fewer rewards in recent history (**Figure 2B,D,E,G,I,J**), replicating a previous finding on a similar task^27^. Reasoning that ELA might alter how dopamine signals track environmental richness, we identified a window in which dopamine was most strongly correlated with reward history across both groups (see Methods and for details). We then extracted the mean dLight signal during this window (1000-1750 ms after lever press; see **Figure 2B-C** inset) for further analyses.

**Figure 2:**
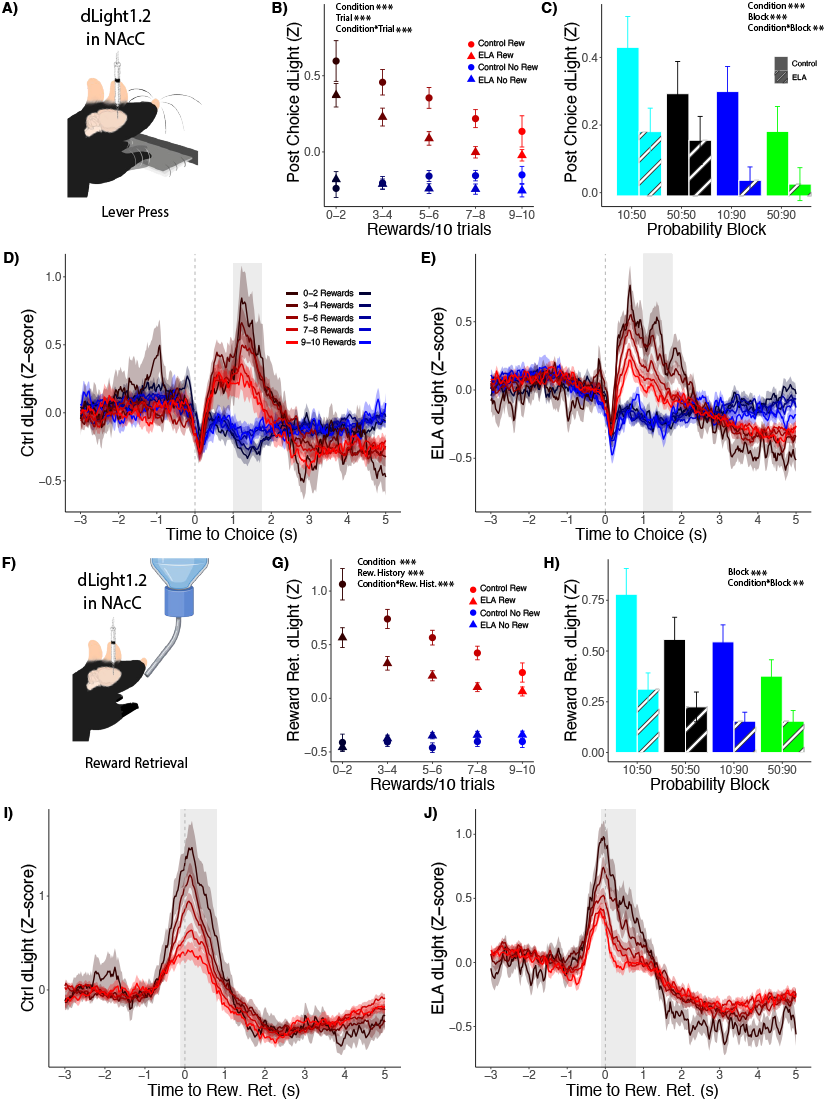
dLight imaging of dopamine release at lever press and reward retrieval. **(A,F)** Animals play a probabilistic reward task with simultaneous *in vivo* dopamine fiber photometry recordings. **(B-C)** Summaries of mean dopamine release during the analysis window shows a clear effect of reward history **(B)** and block **(C)** with more dopamine when there were fewer rewards in recent history and in poor versus rich reward blocks. **(D-E)** dLight signal reflects a phasic increase in dopamine release for rewarded, and a phasic decrease in dopamine release for unrewarded lever presses for Control **(D)**, and ELA **(E)** animals. The magnitude of dopamine release at the time of lever press depends on trial history for both groups with higher magnitude release when there were fewer rewards in the last 10 trials. Grey vertical bars indicate an analysis window from 1,000 to 1750 ms. Window timing was chosen based on the timepoints with the most reliable correlation between dopamine release and reward history for both groups. **(G-J)** dLight signal time-locked to nose-poke for reward reveal dopamine ramping to the point of reward retrieval, and larger magnitude dopamine release when there were fewer rewards on the last 10 trials and in poor versus rich reward blocks. **(G-H)** Grey vertical bars indicate an analysis window from -100 to 850 ms with respect to nose-poke, based on timepoints with the strongest correlation between dLight signal and reward history. *All error bars and bands are +/- SEM*.

### Early life adversity reduces the effect of reward on dopamine release

An analysis of dLight signal in response to rewarded lever presses indicates that ELA alters how dopamine signals reward. A trial-wise mixed effects regression of dLight signal on trial outcome, reward history, group, and the interaction of group and outcome reveals that dopamine release is greater following rewards versus punishments (β = 0.279, p < 2.00 × 10^-16^), and that the effect is smaller on trials in which there were more rewards in the last 10 trials (β = -0.0645, p = 1.81 × 10^-14^). The effect of reward is also smaller among ELA animals versus control animals (β = -0.0795, p = 8.26 × 10^-13^), consistent with human work suggesting that exposure to ELA diminishes the capacity for striatal dopamine signaling^1,36,37^. To minimize confounds posed by between-animal differences in signal intensity, we focus our subsequent analyses on group differences in within-animal effects of dLight signal. For example, we find a reward by reward history interaction such that dLight signaling to reward is negatively related to a history of rewards over the past 10 trials (β = -0.0949, p < 2.00 × 10^-16^). Consistent with slower adaptation of reward expectations in animals exposed to ELA, this reward by reward history interaction is smaller in ELA animals (β = 0.0624, p = 2.33 × 10^-8^, **Figure 2B**).

Mirroring the effect of local trial history, dLight signal is highest in the poor block (where rewards are more scarce) and lowest in the rich block and easy discrimination blocks (where rewards are more plentiful; **Figure 2C**). This pattern is expected under a standard reward prediction error account where dopamine signals of reward prediction errors should be larger when animals are least likely to expect reward. There is also evidence that the adaptation of dopamine signals to rewards across blocks is slower among ELA animals versus controls. In a mixed-effects regression of trial-wise dLight signal on probability block, condition, and their interaction, controlling for reward history, there is significantly higher dLight signal to reward versus punishment in the poor compared to the rich block (β = 0.115, p = 1.80 × 10^-6^) and also in the rich block versus the easy discrimination block (β = 0.0625, p = 7.80× 10^-3^). Importantly, the effect of the rich versus poor block is weaker among the ELA animals (interaction of block type, reward, and condition: β = -0.0813, p = 0.0104), and this is also true in the contrast of the easy discrimination versus the rich block (β = -0.0706, p = 0.0237). Paralleling the behavioral results, these data suggest that ELA animals’ dopamine dynamics are also slower to adapt to changes in block, again supporting a hypothesis of slowed updating of reward expectations among ELA animals.

Replicating prior work^27^ we find that dLight signal ramps up leading up to reward retrieval (**Figure 2I-J**). As with dLight signal following lever press, there is also a parametric effect of reward history on total dLight signal to reward (**Figure 2G,I,J**). A trial-wise mixed effects regression of dLight signal to reward retrieval on reward history, group, and the interaction of group and trial outcome reveals that dLight signal is smaller on trials in which there were more rewards in the last 10 trials (β = -0.232, p < 2.00 × 10^-16^), but the effect of reward history is again blunted in ELA animals (β = 0.125, p = 1.89 × 10^-13^; **Figure 2G**). A mixed effects regression of dLight signal during reward retrieval on probability block, condition, and their interaction, controlling for reward history over the last 10 trials, reveals that dLight signal release is smaller on rich (β = -0.229, p = × 10^-9^) and easy blocks compared with the poor block (β = -0.224, p = 1.66 × 10^-9^). Also, both effects are attenuated among ELA versus control animals (β = 0.222, p = 3.07 × 10^-6^; β = 0.209, p = 1.24 × 10^-5^, respectively; **Figure 2H**).

Collectively, these data suggest that dopamine dynamics reflect reward expectations, adapting to local environmental richness either in terms of cumulative reward history or block, consistent with the reward prediction error hypothesis. Importantly, in both cases, adaptation of expectations is weaker among ELA animals versus controls.

### Dopamine predicts aberrant reward learning in ELA mice

We next tested how learning is shaped by dopamine release. To establish an effect of dopamine release on learning, we looked at whether all animals are more likely to repeat a rewarded action when DA release is higher (**Figure 3B**). In a hierarchical logistic regression of choice repetition, we find effects of condition, reward history and dLight signal following the repetition of the prior choice. Specifically, all animals are more likely to repeat choices when dLight signal at the time of press (1000-1750 ms after lever press) on the prior trial was higher (β = 0.128, p = 0.0188; **Figure 3B**), and when animals have received more rewards in the last 10 trials (β = 0.156, p = 4.92 × 10^-13^; **Figure 3D**).

**Figure 3:**
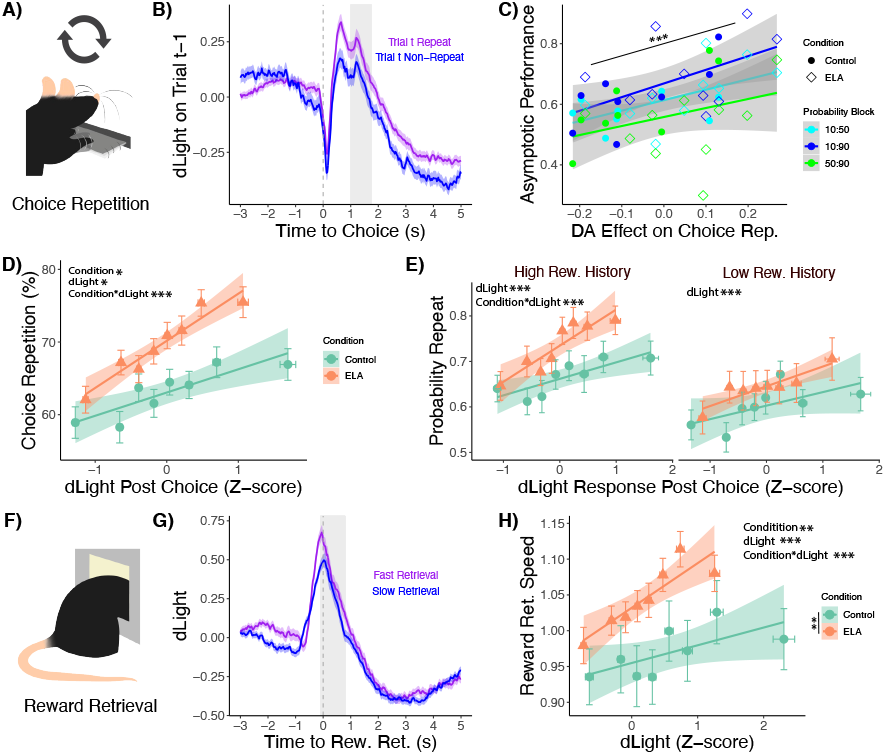
dLight signal reveals that dopamine shapes learning and reward vigor. **(A,F)** Schematic of repeated choice and reward retrieval, respectively **(B)** dLight signal time-locked to lever press indicates that dopamine release to reward on trial T-1 was higher when animals repeated that choice on trial T. This inference was supported by a mixed effects logistic regression revealing that trial-wise dopamine release on trial T-1 predicts an increased likelihood of repeating rewarded choices on trial T. **(C)** The mouse-level random effects of dopamine release on choice repetition correlate positively with asymptotic optimal choice, indicating that animals who show a stronger coupling between dopamine release and choice repetition also perform better, asymptotically. **(D)** The logistic regression further indicates that the effect of dopamine on choice repetition is higher for ELA versus control animals. The plot shows the mean dLight signal on rewarded trials, and mean choice repetition rate following rewarded trials broken out into eight within-animal bins for dLight magnitudes. **(E)** The effect of dopamine and the interaction with group are both stronger in rich (> 5 rewards on the past 10 trials) versus poor trials (< 5 rewards). **(G)** dLight signal time-locked to reward retrieval indicate that dopamine release leading up to nose-poke is higher when animals retrieve rewards faster than their own median retrieval times (and lower when they responded slower). **(H)** Median speed (inverse latency) to retrieve rewards increases with mean dopamine release, time-locked to reward retrieval. All regression plots include 95% confidence intervals on the maximum likelihood regression line. All error bars are +/- SEMs.

Dopamine also has a greater effect on choice repetition and reward history in ELA animals. Specifically, interactions with condition indicate that the effect of dLight signal on repeating choice is larger among ELA animals versus controls (β = 0.185, p = 0.0131), and the effect of reward history is also greater (β = 0.137, p = 4.68 × 10^-6^; **Figure 3D**). That is, for a given unit of dLight signal or for a given number of rewards on the last ten trials, ELA animals are even more likely to repeat a choice as compared with controls. Furthermore, a three-way interaction indicates that the effect of dLight signal on repeating an action is greater among ELA mice in richer environments (when there were more rewards in the last ten trials: β = 0.114, p = 1.72 × 10^-4^; **Figure 3E**).

These results indicate that dopamine release facilitates reinforcement learning, making animals more likely to repeat actions associated with dopamine release. They further indicate that the effect of dopamine release on choice repetition is stronger among mice exposed to ELA. An important question is whether this effect of dopamine on choice repetition impacts goal pursuit. We thus tested whether asymptotic optimal choice is related to the coupling between dopamine release and choice repetition.

To test whether the link between dopamine release and choice repetition is a good proxy for adaptive reinforcement learning, we tested whether the coupling between dLight signal and choice repetition predicted asymptotic optimal choice (**Figure 3C**). Specifically, we extracted the random effects of our trial-wise logistic regression which hierarchically estimates the effect of dLight signal on choice repetition for each mouse and correlated these random effects with asymptotic performance. A hierarchical linear regression of asymptotic performance by block on the extracted random effects reveals that animals with a stronger effect of dLight signal on choice repetition also choose the optimal lever at a higher rate, asymptotically (β = 0.199, p = 1.73 × 10^-4^). This relationship is furthermore independently significant for asymptotic performance in the rich (β = 0.526, p = 0.0282), poor (β = 0.447, p = 0.0492), and easy discrimination blocks (β = 0.533, p = 0.019). These results indicate that a stronger coupling of dopamine signaling and choice repetition predicts more optimal learning.

In summary, one unit of dLight signal produces a stronger learning effect, increasing choice repetition in ELA as compared with control animals. Also, stronger coupling between dLight signal and choice repetition positively predicts asymptotic optimal choice rates. However, ELA animals perform more poorly, on average, in the full sample. Reconciling the observations that each unit of dLight signal produces stronger learning in ELA animals, and that ELA animals also tend to perform worse, on average, requires that the mechanisms driving dopamine release must be blunted in ELA. That is, the effect of dopamine signaling on learning may be stronger in ELA animals, so ELA animals must have weaker dopamine signaling in the first place. Again, we recognize that because dLight signal is not an absolute measure of dopamine, it is possible that ELA animals have lower dopamine concentrations and thus require relatively larger dopamine signaling to achieve the same outcome. The possibility that ELA animals have less dopamine is different from the possibility that ELA animals simply do not regulate their dopamine signaling as much, as a function of learning. We consider this latter possibility next.

Weaker dopamine signaling among ELA animals could be accounted for by blunted mechanisms driving learning about reward contingencies. As ELA animals learn more slowly over blocks (**Figure 1A**), we hypothesized that dopamine signals also adapt more slowly in ELA animals. To test this hypothesis, we examined the effect of reward history on dLight signaling. According to the reward prediction error hypothesis, trial-wise dopamine release should decrease as animals have more experiences with reward predicting cues. As experiences accumulate, animals should make better predictions and release less dopamine at the time of the cue. That is, if the evaluative mechanisms driving dopamine release are blunted, then dopamine release to reward will depend less on reward history. Indeed, as noted, the effect of reward history (rewards in the last ten trials) on dLight signal to rewarded presses is attenuated in ELA versus control mice (**Figures 2I-J and 4D**).

One consequence of this three-way interaction is that ELA animals release relatively less dopamine when rewards are rare and relatively more dopamine when rewards are common. This pattern conceptually aligns with the hypothesis that striatal dopamine signaling shapes behavioral opportunism: weaker effects of dopamine on learning and responding when rewards are infrequent and distant in poor environments, and stronger effects of dopamine in locally rich environments (e.g., when rewards are immediately available).

Controlling for the overall amount of dLight signal as a proxy of dopamine release in response to rewards, ELA animals have a stronger effect of dLight signal on choice repetition. However, because this coupling positively predicts better asymptotic performance across all animals, we infer that ELA is associated with slowed updating in the mechanisms regulating dopamine release. One interpretation, from the perspective of an actor-critic architecture and its biological variants^39,40^ is that ELA animals have a more sensitive actor (adjusting choices more strongly as a function of dopamine), but a less sensitive critic (updating predictions more slowly), thus driving less robust dopamine signaling in the first place. We next test predictions regarding learning algorithms in

### Learning model reveals that ELA slows reward predication error updating

Slowed learning due to blunted adaptation of dopamine signaling with reward history implies that that ELA also slows trial-by-trial adaptation of prediction errors. To test this prediction, we fit a reinforcement learning model to choice behavior in the full set of animals, hierarchically, with sessions nested within in mice. We fit a diverse set of models with differing assumptions about underlying algorithms. The best fitting model (the “perseveration model”) is a learning model with delta-rule updating, and a bias to repeat prior choices (to perseverate) which decays over trials (as index by a perseveration decay term). To confirm the quality of the model fit, we conducted model validation checks by simulating data from the fitted model and comparing to real data. These checks reveal that the perseveration model captures key features of actual performance. Namely, it 1) recapitulates trial, block, and condition effects on optimal choice rates (**Figure 4A**) and 2) captures the effects of trial outcome and reward history on the likelihood of choice repetition (**Figure 4B-C**).

**Figure 4:**
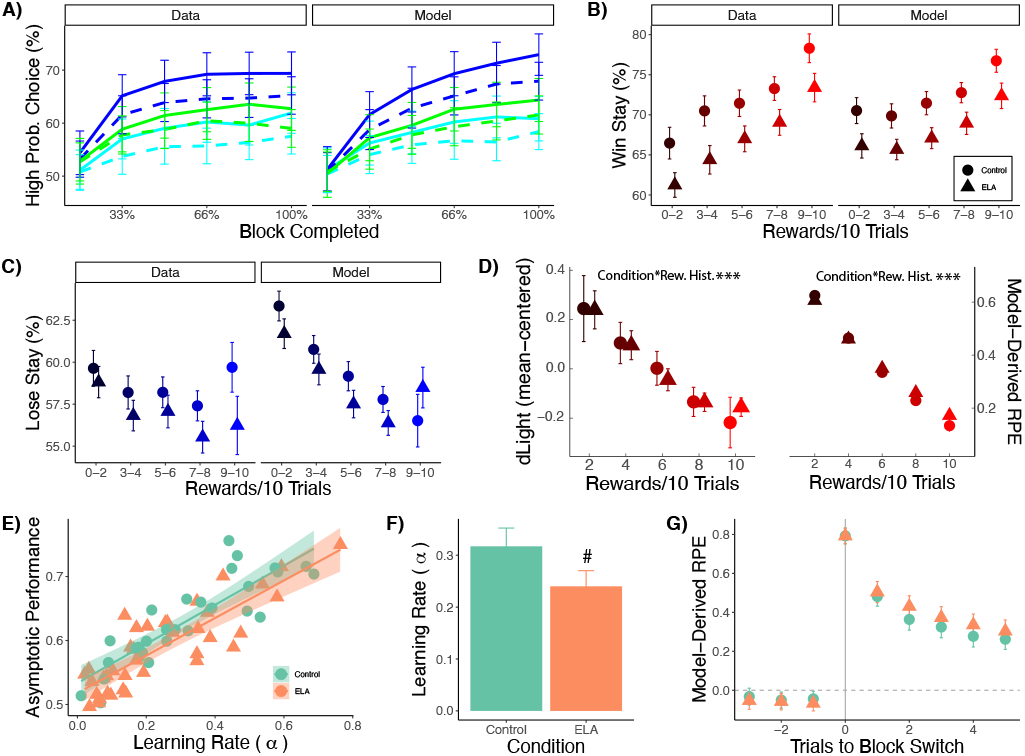
Reinforcement learning model of behavior. The best fitting model, across a range of Rescorla-Wagner type reinforcement learning models is a delta rule update model with a perseveration term biasing repetition of choices from prior trials which decays across trials. **(A-C)** Model validation includes comparing real and simulated data indicating that the perseveration model captures all key features of choice data including the effects of block, trial number within block, reward history, and group on optimal choice rates and choice repetition following re-warded and un-rewarded trials. **(D-E)** Both dLight **(D)** signal and model-derived reward prediction errors **(E)** show the same pattern of decreasing with increasing numbers of rewards on prior trials. Moreover, both show a blunted effect of reward history for ELA versus control animals. **(F)** Model learning rates account for a large fraction of individual differences in asymptotic optimal choice rates, explaining a considerable portion of the group differences in performance. **(G)** Learning rate parameter estimates averaged across sessions for each animal were numerically, trending lower for the ELA versus control animals. **(H)** Slowed learning rates are reflected in slowed adaptation of reward prediction errors with reward experiences in ELA versus control animals.

After confirming that the winning model is also a good model of behavior, we next tested the prediction that trial-by-trial prediction errors reflect slower updating among ELA versus control mice. Specifically, we extracted trial-wise reward prediction errors from simulated data and tested whether RPEs are sensitive to reward, reward history, and condition. A mixed-effects regression of RPEs reveals that – just like dLight signals – model-derived RPEs are larger following rewards versus punishments (β = 1.68, p < 2.00 × 10^-16^) and the effect of reward is smaller when there are more rewards over the past 10 trials (β = -0.232, p < 2.00 × 10^-16^). Importantly, there is also a three-way interaction indicating that the interaction between reward history and reward is blunted in ELA animals versus controls (β = 0.00698, p = 0.027). This threeway interaction (**Figure 4E**) provides computational evidence that ELA animals update their reward expectations and therefore reward prediction errors more slowly than their control counterparts.

The parallel between dopamine release in response to rewarded presses in the dLight mice and the model-derived reward prediction errors from the behavior of the full sample, supports the interpretation that phasic dopamine release conveys reward prediction error signals driving learning. This parallel also implies that mechanisms driving adaptation of reward expectation are slowed in animals exposed to ELA, helping to account for their lower performance on the bandit task.

### Dopamine shapes aberrant vigor in ELA mice

While behavioral opportunism is characterized by slower updating about reward expectations, it is also characterized by more vigorous action when rewards are immediately available. We hypothesize that faster reward retrieval among ELA animals is also driven by striatal dopamine signaling – albeit via an effect of dopamine signaling on the performance rather than the learning of actions^27,35,39,41^. Indeed, a median split of trials based on within-mouse reaction times reveals stronger dopamine signaling on fast relative to slow response trials (**Figure 3G**). To test the hypothesis that dopamine drives relatively more vigor for reward retrieval among ELA animals, we regressed trial-wise reward retrieval speed (inverse latency) on condition, and dLight signal during the reward retrieval window (−100 ms before to 800 ms after nose-poke), controlling for reward history on the last 10 trials (**Figure 3H**). We find that, as predicted, the effect of dLight signal on vigor differs by group (β = 0.0513, p = 0.00944). The sign of the interaction indicates that each unit of dopamine released leading up to reward produces faster reward retrieval times in ELA versus control animals. Thus, dopamine signaling drives a group difference in greater vigor among ELA animals in proximity to reward.

### Exposure to kicking accounts for variance in blunted task performance beyond ELA

Slower updating of reward expectations is a rational strategy when the world is perceived as more unpredictable. We hypothesized that slowed learning in behavioral opportunism arises from early experiences of unpredictability in both the quantity and quality of early life care. Prior work revealed that animals in the limited and bedding nesting paradigm (ELA) experience inconsistency in maternal care as indexed by higher rates of maternal nest entries/exits^42–45^ (**Figure 5C**) and exposure to maternal kicking^43^; **Figure 5A-B**). Furthermore, there was evidence that exposure to maternal kicking subsequently altered how animals weighed costs and benefits in adulthood. Thus, we predicted that nest entries and kicking correlate with worse performance on the probabilistic band task, over and above condition. Independent regressions, controlling for condition, reveal that both maternal kicking and nest entries are reliably related to worse rates of optimal choice. Specifically, the frequency of kicking and the nest entries/exits accounts for a large fraction of between-individual variance in asymptotic optimal choice (**Figure 5C-D**). Specifically, animals which were kicked more (β = - 0.331, p = 0.0137), or experienced a caregiver who was less consistent in their caregiving (at trend-level: β = -0.263, p = 0.0616) chose the optimal lever at lower rates. Maternal kicking in ELA animals predicted reliably worse asymptotic optimal choice rates (β = -0.251, p = 0.00961), but kicking did not predict performance in the control group (β = 0.166, p = 0.183). However, nest entries are reliably predicted worse asymptotic optimal choice rates in the control group (β = -0.633, p = 1.17 × 10^-8^), but not in the ELA group (β = 0.131, p = 0.256). Taken together, the ELA paradigm induces slowed reward learning in adulthood, but even more important for performance are individual differences in the quantity and quality of early life care.

**Figure 5:**
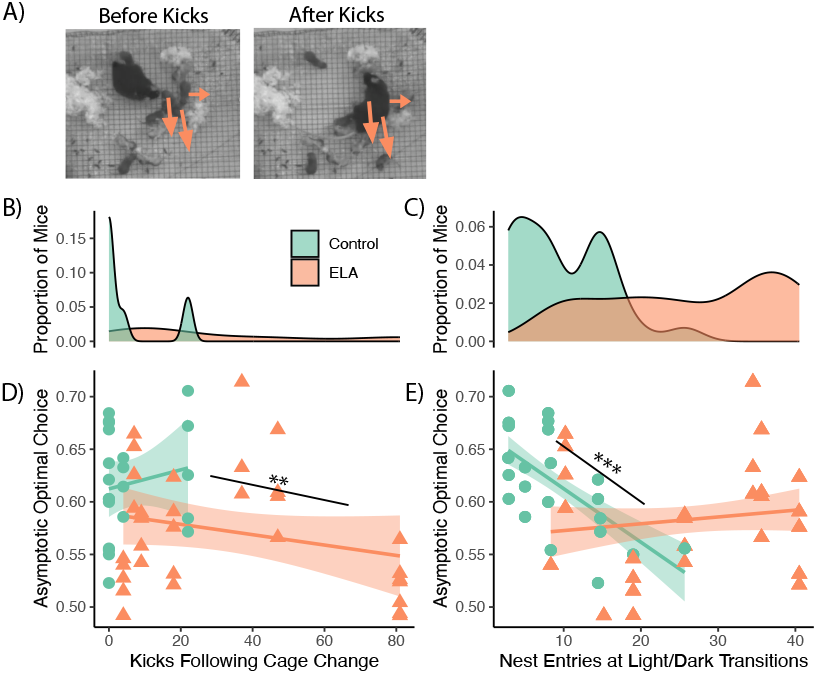
Early life experience predicts asymptotic optimal choice. Mice reared in the ELA condition experience diminished consistency and quality of maternal care. **(A)** Kicking refers to mom displacing pups from nest with a rough front or hind less kick. Specifically, the ELA pups experience mom **(B)** leaving and entering the nest more frequently compared with Controls and **(C)** exhibiting abuse-like pup kicking, which is almost non-existent among Control dams. Both quantities were measured during one hour of home cage video recordings from post-natal day 4. Nest entries are averaged across two timepoints in the day, at 6:00 and 18:00. **(D-E)** Scatterplots show that asymptotic optimal choice across the easy discrimination, rich, and poor blocks depend on these factors beyond group membership. **(D)** Higher maternal kick rates experienced in early life correlate with lower asymptotic performance in the bandit task across all adult animals. **(E)** A higher rate of maternal nest entries/exits does not correlate with asymptotic optimal choice across all animals, but there is a strong relationship of more nest entries/exits correlating with worse optimal choice for the Control animals only. ELA animals exhibit lower asymptotic optimal choices overall, but there is no further correlation with nest entries/exits. Bands show 95% confidence intervals on the least squares regression

## Discussion

In this study, we investigated whether early life adversity (ELA) impacts reward learning and decision making across the lifespan by altering striatal dopamine signaling. We hypothesized that animals adapt to ELA by developing a strategy of behavioral opportunism. Specifically, that animals update their reward expectations more slowly, and conserve their vigor for uncertain rewards, but act more vigorously when reward is immediately available (a rational strategy in the face of uncertainty). Prior work on the effects of ELA indeed has indentified these patterns in adult humans^4,7,9,12–14,17,46^, and also that they may underlie ELA individuals’ increased risk for pscyhopathologies such as substance use disorder and depression^17,47,48^.

To test our hypotheses, we evaluated the effects of limited bedding and nesting or control rearing conditions on a reward learning and decision-making task that systematically manipulates reward statistics and environmental richness in adult mice. We used dLight fiber photometry to evaluate striatal dopamine dynamics during the task in a subset of mice.

Our groups differed in overall task performance. As predicted, ELA animals learn at a slower rate and choose optimal (reward maximizing) levers less often, asymptotically. However, when rewards are immediately available, the same ELA animals that learn more slowly, also retrieve rewards more vigorously. These behavioral findings support our prediction that ELA predicts a strategy of behavioral opportunism that persists into adulthood. We further hypothesized that ELA-induced disruptions to striatal dopamine signaling^1,36,37^ underlie the development of behavioral opportunism. To test this hypothesis, we examined whether there were relationships between aberrant dopamine signaling, disrupted learning, and increase vigor for available rewards.

### Dopamine predicts blunted learning in ELA animals

What neural mechanisms underlie ELA animals’ slowed reward learning? One simple explanation is that ELA mice release less dopamine overall, thus inducing less plasticity. Consistent with this account, we find lower striatal dLight signals in ELA mice in response to rewards (cf. **Figure 2A-B,H-I**). Thus, ELA mice may be slower to learn because they release less dopamine in response to rewards, blunting their reinforcing effects. While we z-scored dLight signals within mice, thereby putting all animals on the same scale, we acknowledge that differences in overall signal intensity unrelated to striatal dopamine release may account for some fraction of this group difference. Thus, we restricted our inferences, more conservatively, by examining group differences in terms of within-animal effects of striatal dopamine signals on behavior, and how these adapted with learning.

We first analyzed the effect of dLight signal as a proxy for dopamine release in response to a rewarded press on the probability of repeating a choice. As expected, when dLight signal is higher, all animals are more likely to repeat rewarded choices on the next trial, and these effects are stronger in rich versus poor environments (when there were more rewards in recent trials). These results indicate that dopamine release, as measured by dLight signaling, promotes adaptive learning for reward pursuit. We further found that animals with a stronger coupling between dLight singaling and choice repetition make optimal choices at a higher rate, asymptotically. This indicates that the link between dLight signal to press and choice repetition is a good index of the effects of striatal dopamine release on adaptive reward learning.

Surprisingly, we found that ELA animals, despite being slower learners overall, show a stronger link between dLight signal and choice repetition. That is, for a given unit of dLight signal, ELA animals are more likely to repeat rewarded choices relative to control animals. Taken together with the observation that ELA animals generally learn more slowly and perform worse, we infer that the mechanisms which signal reward prediction errors and update reward expectations must be blunted in ELA animals. We tested the specific prediction that reward expectations are updated more slowly, according to reward history, among ELA animals relative to controls. Indeed, we found that while striatal dLight signal to reward decreases with greater reward in recent history, this signature of reward prediction error signaling reflects slower learning / adaptation among ELA mice.

The effect of ELA strengthening the link between dLight signal and choice repetition is also consistent with the interpretation that ELA induces behavioral opportunism by enhancing exploitation of more proximal rewards. Indeed, computational models of dopamine and action selection in the striatum suggest that one impact of dopamine on choices is to more reliably seek more rewarding options – a strategy which is adaptive for exploitation^39^. Thus, what counts as rewarding, for animals exposed to ELA, should have a greater effect on subsequent choice and this would be driven by dopamine-mediated plasticity. We speculate that this ELA effect may have to do with changes in receptor density in response to dopamine signaling.

Computational modeling of behavior in the full sample of mice reinforced the inference that ELA animals learn more slowly. Just as ELA slows the updating of striatal dopamine signals as a function of reward history, ELA reward prediction errors (generated by the model fit to ELA mice behavior) also update more slowly as a function of reward history. The parallel in the empirical and simulated results buttresses the inference that aberrant plasticity and striatal dopamine underlies slowed learning in ELA.

One explanation for aberrant dopamine-mediated plasticity is that ELA animals have more uncertainty about reward statistics which could blunt reward prediction errors especially when rewards are statistically rare (e.g., in poor environments). Importantly, ELA animals are more sensitive to the distinction between rich and poor environments. Namely, while both ELA and control animals show a heightened propensity to repeat rewarded actions with greater dopamine release, and this effect becomes stronger in rich versus poor environments, such effects are stronger among ELA mice. This pattern is consistent with an interpretation of uncertainty-based blunting of reward prediction errors among ELA mice, especially in poor environments.

Another explanation for blunted dopamine release is structural in nature. ELA animals may synthesize dopamine at a lower rate or have blunted vesicle transport or a lower prevalence of dopamine terminals in the striatum. In fact, human work has suggested lower dopamine synthesis capacity among individuals exposed to adversity during development^37^. Finally, we also considered that slowed learning might reflect alterations in striatal dopamine receptor density. For example, more pre-synaptic D2 receptors may enhance auto-receptor function, thus curtailing dopamine release in response to rewarding events. Future work should assess the relative contributions of structural differences or adaptive dopamine signaling to slowed learning among ELA mice.

### Dopamine predicts ELA animals’ faster reward retrieval

In contrast to the observation that ELA animals are slower to learn, when reward is immediately available, ELA animals show greater sensitivity - retrieving rewards more vigorously. This group difference does not reflect differences in overall vigor. In fact, while ELA mice were slower to make their choices, they were faster to act when reward was immediately available. Dopamine appears to underlie this effect as well: dLight signal during reward retrieval predicts greater vigor among ELA mice versus controls. Prior work in this same task has shown that the intensity of dLight ramping during reward retrieval is related to animals’ willingness to work for reward, producing greater vigor^27^. Critically, this effect is amplified among animals exposed to ELA.

One explanation for the heightened effect of reward and dLight signal on vigor is that when a discriminative cue indicates reward availability, the uncertainty that ELA animals face is resolved, permitting more robust effects on reward retrieval. Notably, the effect of dLight signal on vigor is enhanced in reward rich environments where animals may have the least uncertainty about the probability of reward (**Figure 3**). A core premise of behavioral opportunism is that animals withhold their effort until they are certain it will pay off. Given the intense and chronic uncertainty faced by animals reared in adverse early environments^43^, this would seem a rational adaptation. Heightened sensitivity to reward uncertainty, among ELA animals, may generally blunt dopamine release to reward predictive cues. ELA animals are also likely to be more uncertain about reward availability, in general, given that they are slower to learn. Indeed, ELA animals are slower to act at the time of choice, which may reflect greater uncertainty. While the experience of reward tends to speed choices on the next trial, this is less true for ELA animals (**Figure 1**). Thus, the same experiences of reward may confer less certainty for ELA animals than they do for control animals.

### Early life experiences account for variance in task performance beyond group

A key prediction of our prior work^43^ was that the degree of uncertainty in both the quantity and quality of maternal care is related to adult outcomes. We tested this prediction in relation to rates of asymptotic optimal choice. We found that ELA was associated with less consistent maternal care (e.g. high rates of maternal nest entries / exits) as well as poorer quality care (e.g. higher rates of maternal kicks), replicating our prior study. Critically, these outcomes were related to worse asymptotic performance across all mice and explained variance in performance over and above group differences. Thus, animals who experience more variable maternal care, and also abuse-like behavior are worse at probabilistic reward learning.

### Limitations

Our work has several limitations. We imaged dLight signal exclusively in the ventral striatum which may have different implications for behavior as compared to dopamine signaling in the dorsal striatum. While the former has classically been linked with state evaluation (the critic), the latter has been associated with learning action policies (the actor)^40,49^. Thus, dorsal striatal dopamine may bear a closer relationship with action learning and execution then the dynamics we evaluated in the ventral striatum. Another limitation is that we used dLight fiber photometry, which is subject to bleaching and may be susceptible to differences in florescence across animals. Future work using recent advances in cyclic volumetry may provide better-calibrated measures of dopamine dynamics.

The corollary hypothesis of behavioral opportunism is that ELA animals conserve their energy when reward is not immediately available to avoid incurring effort costs. While our behavioral paradigm manipulated the potential benefits of actions by varying reward probabilities, we did not manipulate potential costs. Future experiments should incorporate variations in punishment time outs or physical effort requirements to explore the effects of early life adversity on sensitivity in these cost domains.

## Conclusion

Our results support the hypothesis that animals exposed to ELA adapt by developing a strategy of *behavioral opportunism*. Moreover, this strategy is partly mediated by aberrant striatal dopamine function. We speculate that aberrant striatal dopamine signaling and concomitant *behavioral opportunism* lay the foundation for diverse forms of reward-related psychopathologies and susceptibility to substance use disorders in humans exposed to ELA.

## Acknowledgements

We thank Andrew Westbrook, Gabriela Manzano Nieves, Rebecca Burwell, and Robert Reuter for their invaluable discussions. This work was supported by funding from the National Institute of Mental Health on Drug Abuse (NIDA) grant F31DA053088 to M.G and. National Institute of Mental Health (NIMH) grant R01MH115049 to K.G.B.

## Author contributions

M.G.: conceptualization, investigation, methodology, formal analysis, visualization, writing—original draft. A.A.H.: conceptualization, methodology, investigation, formal analysis, visualization, writing—review and editing. A.J.: conceptualization, methodology, formal analysis, visualization, writing—review and editing. T.P.: investigation. D.O.: investigation, visualization, writing—review and editing. C.D.: investigation, writing—review and editing. . J.B.: investigation, writing—review and editing. A.I.M.: investigation. M.J.F.: conceptualization, supervision, formal analysis, writing— reviewing and editing. C.I.M.: conceptualization, supervision, formal analysis, writing—reviewing and editing. K.G.B.: conceptualization, supervision, formal analysis, writing—reviewing and editing.

## Materials and Methods

### Animals and Housing

All animals (including 38 control mice and 39 early life adversity mice) procedures and maintenance complied with Brown University’s Institutional Animal Care and Use Committee and in accordance with the National Institutes of Health Guide for the Care and Use of Laboratory Animals. C57BL/6 mice were bred in house and maintained on a 12:12 light/dark cycle (lights on at 6:30, off at 18:30) with *ad libitum* access to food and water. Mice were housed in 31 (l) x 12 (w) x 14 (h) cm cages with bedding and a 4x4 cm cotton nestlet. Pups were weaned, segregated according to sex and pair housed at postnatal day 21.

### Limited Bedding and Nesting (LBN) Manipulation

Four days following birth of a litter, dams and pups were transferred from their home cage into a 31 (l) x 31 (w) x 14 (h) cm cage with a wire mesh floor with a 4x2 cm cotton nestlet for seven days (P4-P11)^42,43,50^ . Throughout this manipulation, mice continued to have ad libitum access to food and water. Following the limited bedding condition, dams and their pups were returned to normal bedding conditions.

### Continuous Home Cage Video Recording

Mice in either limited bedding and nesting or control conditions were maintained in a specialized housing room in the vivarium, in typical housing cages with a clear plexiglass cage topper with ventilation holes ^43^ . For eight consecutive days (from postnatal day 3-11), cages were continuously recorded in the vivarium from an overhead camera. We scored dam-pup interactions in LBN and control cages across the circadian cycle (0:00, 3:00, 6:00, 12:00, 18:00). All scoring was conducted by four individuals. Inter-rater reliability scores are provided in the supplement (Cohen’s kappa = 0.779).

### Behavioral Task

In adulthood (> PND 65), mice were mildly water restricted to incentivize performance^51^ . All mice were water restricted to approximately 1.5 g of water/day (11% of their weight in water) and maintained at 90% or higher of their free running weight. Training and testing were carried out in Med Associates (size) box with two retractable, ultra-sensitive mouse levers, one mouse dipper with cup (10uL or 1cc) with a head poke port, one house light and two cue lights (above both levers; **Figure 3.1A**). Carnation® evaporated milk was used as a reinforcer. Mouse activity, including lever pressing, head poking and location, were recorded by Noldus Ethovision XT 13 and a subset of mice were recorded by LabView (for simultaneous behavioral and fiber photometry recordings).

In phase one pre-training (60 min), water-restricted mice simultaneously underwent frequency of reinforcer 1 (FR1) and frequency of timed reward 30 s, where mice could either press a lever (randomly one of two levers extended on each trial) for reward or wait thirty seconds for a reward until animals obtained 30 rewards. In phase two (60 min), mice completed FR1 training, pressing one lever for one reward until animals pressed for 30 rewards on two consecutive days. During phase three (90 min), mice were presented with a choice between two levers with reward probabilities of 50%. If mice selected the same lever more than three times in a row, the reward probability of that choice dropped to zero and the other choice increased to 100%. After receiving 60 rewards in each of two consecutive days, animals were deemed ready to advance to the probabilistic bandit task training.

Over the course of a bandit task session (1.5 hours), mice pressed one of two levers during seven blocks of reward probability pairings. In a block, two levers presented all reward probability combinations of 10%, 50% and 90%, except 10:10 and 90:90. Each block consists of one probability pair randomly selected from a uniform distribution for 25-35 trials. At the start of each trial, the two levers are extended, and lever cue lights turn on. The animal had unlimited time to decide between the two levers. If a choice is rewarded, reward is immediately presented via dipper with audible click, and remained available for 5 seconds, followed by an inter trial interval (ITI). If choice is unrewarded, the ITI began immediately. During the ITI (8-15 s, randomly chosen from uniform distribution), the house light turned on for the duration of the time out period. Animals were trained on the task for approximately two weeks, until each individual reached an asymptote in session-by-session performance. After training, animals continued to perform the same task over multiple sessions. Sessions were excluded if animals choose a single lever more than 90% of the session. The median number of sessions included per animal was n = 24.

Optimal choice was defined for each block with a difference in reward probabilities according to the lever with the higher probability. To assess asymptotic performance, we examined left/right choices during the second half of each block, when choices stabilized. For estimating linear mixed effects, we used lmer and glmer from the lme4 package in R. For ex-Gaussian analyses of reaction time distributions, we used the exGauss package in R.

Reaction times to choice were defined as the time between lever presentation and lever press. Reaction times for reward retrieval were defined as the difference between dipper presentation and nose poke. Rewards were available for 5 s after lever presentation – nose pokes after this time were excluded from further analyses.

### dLight 1.2 Fiber Photometry

After three to five months of behavioral training, we injected a subset of randomly-select mice from each group with pAAV5-hSyn-dLight1.2^52^ (AP 1.7, ML .8; DV 3.3 for injection one, 3.5 for injection 2 from bregma) and implanted mice with a head-fixed cannula (ThorLabs Fiber Optic Cannula, Ø1.25 x 6.4 mm Ceramic Ferrule, Ø200 µm Core, 0.39 NA) above the nucleus accumbens core (NAcc; AP 1.7, ML .8, DV 3.4 from bregma). Positioning was verified with post-mortem histology. Mice recovered for at least five days before retraining and were recorded playing the probabilistic bandit task, between 3 and 5 weeks, post-implantation. A final set of (n = 8 control and n = 9 early life adversity) mice were included in the analysis based on having a good signal to noise ratio of dLight response to rewards versus punishments at the time of lever press, yielding a total of (n = 53 control and n = 79 early life adversity) sessions for dLight analyses.

A 0.02 Hz high-pass, 2.5 ripple, 3 coefficient filter IIR filter was applied to dLight data to remove slow drift in the signal across a session. Filtered dLight data were then z-scored for each session. For trial-by-trial comparisons, data were time locked to either trial press or rewarded nose poke.

To extract dLight signal for analysis on each trial, we picked a window in which signal was most strongly correlated with reward history, across both groups. Prior work^27^ has shown that the magnitude of dopamine release in response to a reward cue is sensitive to recent reward history in this task. For our analyses, we determined the window with the strongest correlation between dLight signal amplitude and reward on the last 10 trials, by applying a 750 ms sliding window, and computing the correlation at each step. Visual inspection of p-values indicated that the strongest correlations across both groups, for reward and punishment trials were between 1,000 and 1,750 ms, time-locked to lever press and between -100 and 850 ms, time-locked to reward retrieval. Importantly, the exact position of this analysis window did not matter for our key analyses of group differences in the effect of dopamine on behavior. To confirm this, we re-fitted key mixed-effects regression models from the main text, across a sliding window of timepoints and verified that the specific timing of the analysis window did not change the nature of any reported findings.

Finally, although dLight signals were z-scored within each session so that data could be compared across sessions and animals, we recognize that between-animal or between-session differences in signal fidelity may add noise to our analyses. As such, we more conservatively restricted our analyses to group differences in the within-animal effects of dLight on behavior.

### Reinforcement Learning Model

To estimate learning effects and reward prediction errors within each trial, we fit a diverse set of Rescorla Wagner-type reinforcement learning models using an expectation maximization algorithm in MATLAB (Huys, 2017; for full list of models and model comparison see. We fit our models for each animal separately, across all sessions for each animal. Expectation maximization is a hierarchical fitting procedure that enables simultaneous estimation of parameters at the session and the animal level, assuming that all sessions were produced by a common mechanism (the animal). For between-animal and between-group comparisons, we analyze the session-level parameter estimates and data simulated from session-level parameters.

We performed model comparisons across a range of models according to information criteria (BIC) scores. The best fitting model, for nearly all our mice, was a model that accounted for the high rate of choice repetition (the “perseveration model”) The perseveration model was a Q-learning model consisting of terms for learning rate, inverse temperature, perseveration bias and perseveration decay.

On each trial, the value of an action, *Q*, was updated according to a learning rate, α, and a reward prediction error:

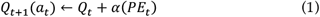

Where the prediction error was calculated from the difference between the actual, *r*, and the expected value of the action:

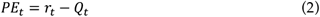

The distinguishing feature of the perseveration model is that it captures animals’ bias, η, to repeat actions that were just taken by increasing the effective action value. This bias falls off with the number of trials since the action was taken, *c*. Note that the trial count for a given action is set to *c* = 1, when an action is taken and increments, at the rate of (0 < δ < 1), for all other trials.

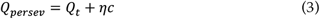

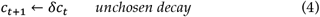

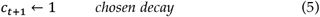

Finally, the probability that an action *i* was selected, was determined according to the softmax function with an inverse temperature, β.

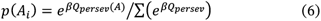

Thus, the perseveration model had four parameters estimated for each session and mouse, α, η, δ, and β. The parameters all had good recovery indicating that they are stable and suitable for inference.

